# Soil nutrient availability alters tree carbon allocation dynamics during drought

**DOI:** 10.1101/2020.08.25.266205

**Authors:** Leonie Schönbeck, Mai-He Li, Marco M. Lehmann, Andreas Rigling, Marcus Schaub, Günter Hoch, Ansgar Kahmen, Arthur Gessler

**Affiliations:** Forest Dynamics, Swiss Federal Institute for Forest, Snow and Landscape Research WSL, Zürcherstrasse 111, 8903 Birmensdorf, Switzerland; Department of Environmental Sciences – Botany, University of Basel, Schönbeinstrasse 6, 4056 Basel, Switzerland; Plant Ecology Research Laboratory, School of Architecture, Civil and Environmental Engineering, EPFL, Lausanne, Switzerland; Institute of Terrestrial Ecosystems – ETH Zürich, Universitätstrasse 16, 8092 Zürich, Switzerland

**Keywords:** ^13^C, drought, fertilization, isotopes, ^15^N, nitrogen allocation, *Pinus sylvestris*, sink strength, carbon allocation

## Abstract

Drought alters allocation patterns of carbon (C) and nutrients in trees and eventually impairs tree functioning. Elevated soil nutrient availability might alter the response of trees to drought. We hypothesize that increased soil nutrient availability stimulates root metabolism and carbon allocation to belowground tissues under drought stress. To test this hypothesis, we subjected three-year-old *Pinus sylvestris* saplings in open-top cambers during two subsequent years to drought using three different water treatments (100%, 20% and 0% plant available water in the soil) and two soil nutrient regimes (ambient and nitrogen-phosphorus-potassium (N-P-K) fertilization corresponding to 5 g N/m^2^/yr) and released drought thereafter. We conducted a ^15^N and ^13^C labelling experiment during the peak of the first-year drought by injecting ^15^N labelled fertilizer in the soil and exposing the tree canopies to ^13^C labelled CO_2_. The abundance of the N and C isotopes in the roots, stem and needles was assessed during the following year. C uptake was slightly lower in drought stressed trees, and extreme drought inhibited largely the N uptake and transport. Carbon allocation to belowground tissues was decreased under drought, but not in combination with fertilization. Our results indicate a potential positive feedback loop, where fertilization improved the metabolism and functioning of the roots, stimulating the source activity and hence C allocation to belowground tissues. This way, soil nutrients compensated for drought-induced loss of root functioning, mitigating drought stress of trees.

## Introduction

Carbon allocation is an important determinant of the C budget of forests and their response to changing environmental conditions. During the growing season, trees actively take up C and allocate it to growth, defense, respiration or storage (Chapin et al. 1990, Körner 2003, Litton et al. 2007). Seasonal fluctuations in C storage pools occur, with refilling of the pools in preparation for early spring growth and depletion of pools when peak growth requires more C than is being assimilated (Oberhuber et al. 2011). 10% to even 50% of the C used for early spring development of new shoots and leaves in deciduous trees may be previously stored C (Hansen 1967, Hansen and Beck 1990). In evergreen coniferous trees, the role of stored C is assumed to be smaller due to photosynthetic activity of the older needles. Drought might affect the use of stored C for the production of new foliage, considering the C limitation and the potential negative effect of drought on phloem transport of stored C. Klein et al. (2014) assumed a close coordination between C supply and demand for the development of new needles in drought exposed *Pinus halepensis* leading to smaller needles rather than to stronger use of carbohydrate storage.

C allocation is generally prioritized to tissues increasing the uptake of limiting resources (Freschet et al. 2018). Changing environmental conditions can thus alter the C allocation strategy of trees. While mild drought has been shown to increase the transport of new assimilates to the roots for the production of larger water absorbing surfaces (Kozlowski and Pallardy 2002), extreme drought events seem to reduce the C supply to roots (Hommel et al. 2016, Salmon et al. 2019), either due to lower water use and photosynthesis, or due to lower belowground sink strength, both leading to reduced phloem transport (Hagedorn et al. 2016, Hesse et al. 2019). The tipping points where a further increase in drought duration or intensity leads to a switch from increased to reduced belowground allocation of C, are however not well described.

Given the fact that the intensity but also the frequency of drought and subsequent rewetting events is predicted to increase in future (Easterling 2000), it is important to better understand the ability of plants to recover from restricted water supply (Feichtinger et al. 2015). Recent studies have shown that trees are able to prioritize C storage over immediate growth during recovery (Sala et al. 2012, Galiano Pérez et al. 2017). Moreover, plant C allocation after drought recovery has been found to be sink-driven, and shortly after rewetting, trees allocate C belowground, probably for restoration of drought-impaired roots (Hagedorn et al. 2016). In general, however, the mechanisms of C allocation that determine the recovery after drought are still far from being resolved.

Not only carbon, but also nutrients are indispensable for growth (Millard and Proe 1992) and survival (Gessler et al. 2017), and the allocation of C and N is tightly related (Gessler et al. 2004, He and Dijkstra 2014). Newly developing leaves and shoots are often supplied by both stored and newly taken up nutrients (Millard et al. 2001). For evergreen trees, remobilization of stored nitrogen (N) can contribute up to 50% of the total N needed for new foliage, and there are indications that lower N storage can reduce the production of new leaves (Millard et al. 2001). Later on, during the growing season, trees rely mostly on root nutrient uptake.

When other resources are not limiting, long-term high N availability is assumed to decrease the root-to-shoot ratio of plants, making them more susceptible to drought events. Moreover, long-term high N availability increases assimilation rates and stomatal conductance and thus leads to greater water loss of plants (Gessler et al. 2017). Drought on the other hand might impair N uptake by the roots, increasing the C:N ratio and inducing nutrient limitation, eventually affecting many processes including stomatal sensitivity to drought and root cell integrity (Gessler et al. 2017). Furthermore, N allocation might be altered by drought, due to the transport of soluble N in the form of amino acids to the roots, to increase tolerance to dehydration (Fotelli et al. 2002).

As ion mobility and nutrient uptake capacity become both impaired when water availability decreases (Kreuzwieser and Gessler 2010), sufficient soil nutrients could increase the available N to the rhizoplane, maintain or even improve general metabolic functions and cell integrity and thus promote a plant’s ability to survive or to recover after a drought (Waring 1987, Gessler et al. 2017). Higher N availability for example might then allow to more efficiently synthesize N-containing osmoprotectants such as proline. These osmoprotectants have positive effects on enzyme and membrane integrity (Ashraf and Foolad 2007) and thus might sustain root metabolism under drought. Severe drought, however, might fully inhibit the uptake of nutrients and their transport from the roots to the leaves independent of the soil nutrient supply. An interaction between drought and soil nutrient availability on tree function is thus likely to occur, due to the tight relation between C and N allocation, but this interaction has not received sufficient attention in research, yet (Gessler et al. 2017).

In this study, we tested how a trees’ C and N allocation during drought and after rewetting is influenced by the availability of nutrients in the soil. For this purpose, we combined ^13^C-CO_2_ pulse labelling of the crowns of three-year-old Scots pine (*P. sylvestris* L) trees with ^15^N-NH_4_NO_3_ labelling. We refer to allocation in two ways: Firstly, we assessed the ^15^N and ^13^C enrichment (^15^N and ^13^C excess based on mg dry biomass) in various plant tissues following isotopic labelling. Secondly, we scaled that enrichment with the total biomass of the respective tissue and calculated the relative ^15^N and ^13^C distribution. We hypothesized that (1) C allocation to the roots increases relative to other tissues under drought but that C allocation to belowground tissues is inhibited if the drought gets too intensive, (2) fertilization results in less C being invested in roots and more in aboveground biomass under optimal water supply, but that with drought, fertilization can improve the C allocation to belowground tissues, especially under more intensive drought, (3) drought stressed trees have a strongly coordinated supply – demand regulation for C and N and thus do not deplete C and N reserves for needle growth early in the season, and (4) rewetting results in enhanced uptake and (re-)allocation of N to the needles when trees grew before under severe water limitation, while at the same time C allocation is prioritized for the restoration of the root system.

## Materials and methods

### Study site

This study was conducted in the model ecosystem facility of the Swiss Federal Research Institute WSL, Birmensdorf, Switzerland (47°21’48’’ N, 8°27’23’’ E, 545 m a.s.l.), which consists of 16 hexagonal open-top chambers (OTC) of 3 m height and a plantable area of 6 m^2^ each. 12 of those chambers were used for this experiment (Supplementary data Figure S1). The roofs were kept closed during the entire experiment to exclude natural precipitation. Belowground, the chambers are divided into two semicircular lysimeters (1.5 m deep) with concrete walls. The lysimeters were filled with a 1 m deep layer of gravel for fast drainage, then a fleece layer that is impermeable for roots but permeable for water, and on top a 40 cm layer calcareous sandy loam soil (Supplementary data Table S1, Kuster et al. 2013). Each lysimeter was planted with 15 three years-old individuals of *Pinus sylvestris* saplings (55.61 cm +/- 5.41 cm height) evenly distributed over the area (approx. 40 cm distance from each other). Temperature and air humidity inside and outside the OTC, as well as soil moisture and soil temperature inside (at 5, 20, 35 cm depth) were automatically monitored (5TM soil moisture and temperature logger, Metergroup, Munich, Germany) (Supplementary data Figure S2). 12 out of the 15 trees were subjected to a defoliation treatment for other purposes (Schönbeck et al. 2020), and three individuals per lysimeter were kept intact. Only these intact ‘control’ trees were considered in this labelling study. Individual tree treatments were given such that trees with similar treatment did not directly neighbor each other, to create an evenly distributed pattern over the area.

### Water and nutrient treatments

The experiment was set up as a split-split plot design. Each chamber was assigned one of three different water regimes as whole-plot treatment (four chambers / replicates per regime). Six sprinklers (1 m high) per lysimeter were evenly distributed, and irrigation was programmed for each lysimeter separately. The amount of water to be applied was controlled by means of the automated soil moisture measurements, where the volumetric water content (VWC) on 20 cm depth was the leading indicator. Field capacity (W100 – 100% water) and wilting point (W0 – achieved by no irrigation at all), the two most extreme regimes, were determined by pF curves, and VWC for the irrigation regimes was adjusted accordingly, allowing for an additional ‘mild drought’, W20 regime, with 20% of the water available compared to W100 (Supplementary data Figure S2). Water treatments started a year after planting to promote proper installation of the plants. The irrigation system, controlling the four soil moisture levels, was in function from April to October in 2016 and from April to mid-July 2017, but not in winter to prevent frost damage. In winter and early spring, watering was done by hand (in W100 and W20) to maintain stable soil water levels. From the 13^th^ of July 2017, all chambers were (re)watered until field capacity was reached (see Supplementary data Figure S2) in order to study the recovery process in the trees.

Twice a year, in April and July, one of the two lysimeters (split-plot) in every OTC was fertilized with liquid fertilizer (Wuxal, Universaldünger, NPK 4:4:3), corresponding to 5 g N/m^2^/year. In April 2016 and in April and July 2017, the fertilizer was applied using 3 L water per lysimeter, and the unfertilized treatment was given 3 L water without nutrients, to prevent differences in water content between fertilization treatments. In July 2016, fertilizer was applied in combination with ^15^N pulse labelling described below. Fertilization significantly increased P and NO_3_ concentration, and the total N pool of plant and soil together (ANOVA) (see soil sampling methods and Supporting Information Figure S3).

### ^13^C and ^15^N pulse labelling

In July 2016 (i.e. in the first year of treatment, Supplementary data Figure S1), a ^15^N pulse labelling experiment was carried out in all irrigation regimes, but only for fertilized plots. Per lysimeter, 34.5 mL of the liquid fertilizer was mixed with 0.85 g ^15^N labelled N (98 atom% ^15^N, in the form of ^15^NH_4_ ^15^NO_3,_ Sigma-Aldrich, Buchs, Switzerland). The amount of ^15^N corresponded to 8% of the total N given, and the total N corresponded to 2.5 g N/m^2^, half of the yearly added amount. 900 mL water was added and the solution was injected with a needle (ø 2 mm) with four lateral holes in the soil, at three different depths (5, 15 and 25 cm), evenly distributed over the planted area (20 cm grid) according to Jesch et al. (2018). The labeling technique allowed (1) to introduce ^15^N into the lysimeter without significantly affecting the actual water and fertilization treatment and (2) to achieve the best possible homogenous spatial and depth distribution of the tracer.

On 10 and 16 August 2016, a ^13^C pulse labelling experiment was conducted. For feasibility, only the W100 and W20 water regimes were selected (4 chambers each). The W20 treatment was chosen above the W0 to ensure photosynthetic activity and thus uptake of CO_2_. Two W100 and two W20 chambers were simultaneously labelled per day. The trees in the chambers were covered with a tall tent of transparent plastic foil. For the labelling application, per chamber, 7.5 g 99% ^13^C sodium bicarbonate (Sigma Aldrich, Buchs, Switzerland) was mixed with 7.5 g standard ^12^C sodium bicarbonate and hydrochloric acid in an airtight sealed beaker outside the chamber to generate the 50% labelled ^13^CO_2_ gas. CO_2_ concentration was measured using a Los Gatos Carbon Dioxide Analyzer (Los Gatos Research, San Jose, USA), which is able to detect both ^12^C-CO_2_ and ^13^C-CO_2_. The labelled gas was pumped into the chamber as soon as the CO_2_ concentration inside reached approx. 300 ppm due to photosynthetic CO_2_ uptake, and was brought to and kept at ∼ 500 ppm for approx. 1.5 hours. Fans inside the chambers ensured an even mixing of the air.

At the time of labelling, there were no drought-induced changes in either C or N concentration in any plant tissue. δ^13^C was on average 1.4 ‰ higher in W20 than W100 trees (Supplementary data Table S2). Moreover, photosynthesis rates and biomass were only slightly but not significantly lower in W20 compared to W100 trees (Schönbeck et al. 2020, Supplementary data Table S3, S4), thus ensuring an active ^13^C uptake of trees in both treatments. Furthermore, this ensures that possible differences observed in label allocation between W20 and W100 would be mainly due to altered strategies induced by drought and nutrient availability.

### Tree harvests and stable isotope analyses

Whole tree harvests took place during the drought treatment in October 2016, July 2017 and, 3 months after rewetting, in November 2017. In general, one complete, living tree per lysimeter (with chamber as replicate, n=4) was sampled including the roots, by excavating the root system until the tree was easily pulled out of the soil. With this technique we could harvest almost the complete rooting system of a tree. The roots of the tree individuals were easily separable as they did not intertwine. In July 2017, after a high mortality in the W0 treatment, we decided to not harvest the surviving trees in W0. Root, stem and needle tissues were separated, dried at 60 °C until stable weight and ground to fine powder. 1 mg (±0.1 mg) of the ground material was weighed in tin capsules and converted to CO_2_ and N_2_ in an elemental analyzer *Euro EA* (Hekatech GmbH, Wegberg, Germany) connected to an Isotope Ratio Mass Spectrometer (IRMS Delta V Advantage, Thermo Scientific, Bremen, Germany) to determine C and N contents and the isotopic compositions. C and N content were assessed as percentage relative to dry weight. Laboratory standards and international standards with known δ^13^C and δ^15^N values were used for calibration of the measurements, resulting in a precision of 0.1‰ for both elements. The isotopic ratios in all samples were expressed in δ notation (‰) relative to the international standard Vienna Pee Dee Belemnite (VPDB) for ^13^C and to N_2_ in air for ^15^N. To calculate the total amount of ^13^C and ^15^N added by pulse-labelling, δ notations were expressed in atom%, as follows:

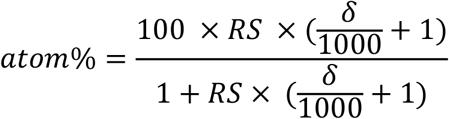

Where RS is the isotope ratio of the international standard (RS: 0.0111802 for ^13^C and N_2_ in air: 0.0036765 for ^15^N) and δ is the δ^13^C and δ^15^N value, respectively. To calculate the excess ^13^C and ^15^N in the plant compartments in µg / g dry biomass, we used

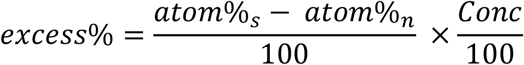

Where atom%_s_ is the atom percentage in the labelled sample, atom%_n_ is the average atom percentage per treatment (water / nutrients) at natural isotope abundance directly before labelling, and Conc is the concentration of C or N in the sample.

Lastly, we calculated the proportion of the total added ^13^C and ^15^N in the plant compartments relative to the total plants’ biomass using

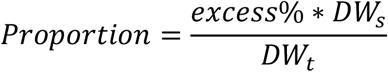

Where proportion is the proportion of ^13^C or ^15^N in a certain plant compartment, DW_s_ is the dry weight of the plant compartment and DW_t_ is the dry weight of the whole tree individual.

### Soil sampling

Soil samples were taken in October 2016, after the first tree harvest. Three soil cores (0-40 cm depth) were taken per lysimeter, evenly distributed over the soil surface to provide a representative soil sample, and the cores were mixed together and sieved. Soil was dried at 40°C, ground to powder, weighed in tin capsules and total N concentration was measured using the EA-IRMS as described above. In addition, 7.5 g dry soil was extracted with 30 ml 1M KCl and filtered through filter paper (Hahnemuehle, Dassel, Germany) into 50 mL PE bottles. NH_4_ concentration in the extract was measured photometrically with flow-injection (FIAS-400) and UV/VIS spectrometer (Lambda 2s, Perkin-Elmer, Schwerzenbach, Switzerland), NO_3_ was measured by colorimetric analysis (Cary-UV50 spectrophotometer), using the absorption of nitrate at a wavelength of 210 nm. Soluble and exchangeable and microbial P were extracted using the method of Hedley (1982), modified by Tiessen and Moir (2007).

### Statistical analysis

Linear mixed effect models were used to test the ^13^C excess, and the proportions within a tree individual, against water treatments and fertilization and their interaction. The individual OTC’s were taken as random factor. ^15^N excess and distribution in the plant was tested for water treatment differences with LMER with individual OTC’s as a random factor. Pairwise differences for both elements were tested with Tukey multiple comparison tests with a Bonferroni correction for multiple testing (package “multcomp” (Hothorn et al. 2019)). Every plant tissue and every harvest time were analyzed separately. All analyses were carried out with R v.3.5.1 (R Core Team, 2019)

## Results

### ^13^C incorporation and distribution affected by drought

Water regimes did not affect the ^13^C excess in any tissue shortly after pulse-labeling during the first year of drought (October 2016), but clear drought effects were observed on ^13^C excess in needles produced in 2017 (N17), roots, and stem in the second year of drought (July 2017) (Figure 1, Supplementary data Table S5). Drought caused an increase in ^13^C in N17 needles compared to the W100 treatment. After rewetting, the effects of the previous water regime were absent (November 2017) (Figure 1, Supplementary data Table S5).

**Figure 1.**
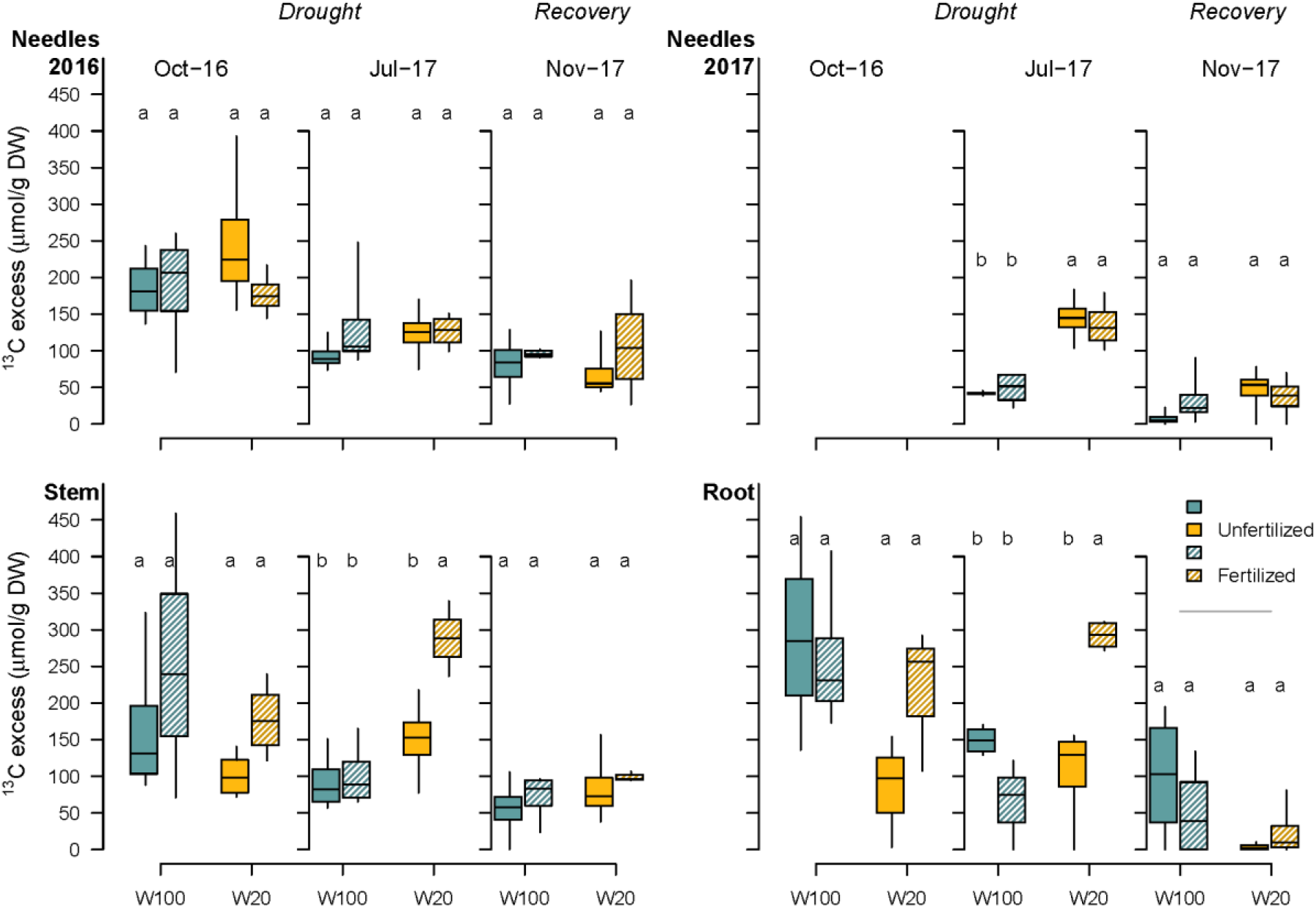
^13^C excess in needles produced in 2017 and 2016, stem and roots in October 2016, July 2017 and November 2017 (three months after rewetting). ^13^C label was applied in August 2016. Water regimes are indicated by colors, fertilization is indicated by shading (solid = unfertilized, pattern = fertilized). Letters indicate significant differences between water and fertilization treatment within every tissue and harvest date. N=4 for every boxplot.

### ^13^C incorporation and distribution affected by combined drought and fertilization

An interaction between the water regime and fertilization was observed in stems and roots, where drought alone decreased the absolute allocation of ^13^C to these organs, while combination of drought with fertilization stimulated it (Figure 1). In line with the absolute ^13^C excess results, only in unfertilized trees, the proportion of ^13^C in the roots decreased from 30% ±4% in W100 to 12% ±5% in W20, whilst fertilized trees had 23% ±4% of the ^13^C in the roots in both W100 and W20 trees (October 2016) (Figure 2). In July 2017, the fertilization effect on root C allocation during drought was even stronger (Figure 2), while in fertilized W100 trees, the proportion of ^13^C in the roots was minimal. An interaction effect of drought and fertilization was also found in the allocation to new grown (i.e. N17) needles. Unfertilized trees allocated relatively more ‘old C’ (^13^C assimilated in 2016) to needle growth in 2017 when affected by drought (8% in W100, 14% in W20), whereas fertilized trees allocated relatively less ‘old’ C in W20 compared to W100 trees (Figure 2). There were no treatment differences in the proportion of ^13^C ending up in the stem, but over time, the proportion of ^13^C that was found in the stem gradually increased in every treatment. After rewetting, the only significant difference was found in needles grown in 2017, where unfertilized W100 trees had the lowest proportion of ^13^C invested in those needles compared to the other treatments. The absence of statistical significance might be due to the high variation in the data, caused by individual variation in recovery. In summary, fertilization stimulated carbon allocation to belowground under drought, whilst ^13^C stayed in aboveground tissues/needles in unfertilized trees under drought.

**Figure 2.**
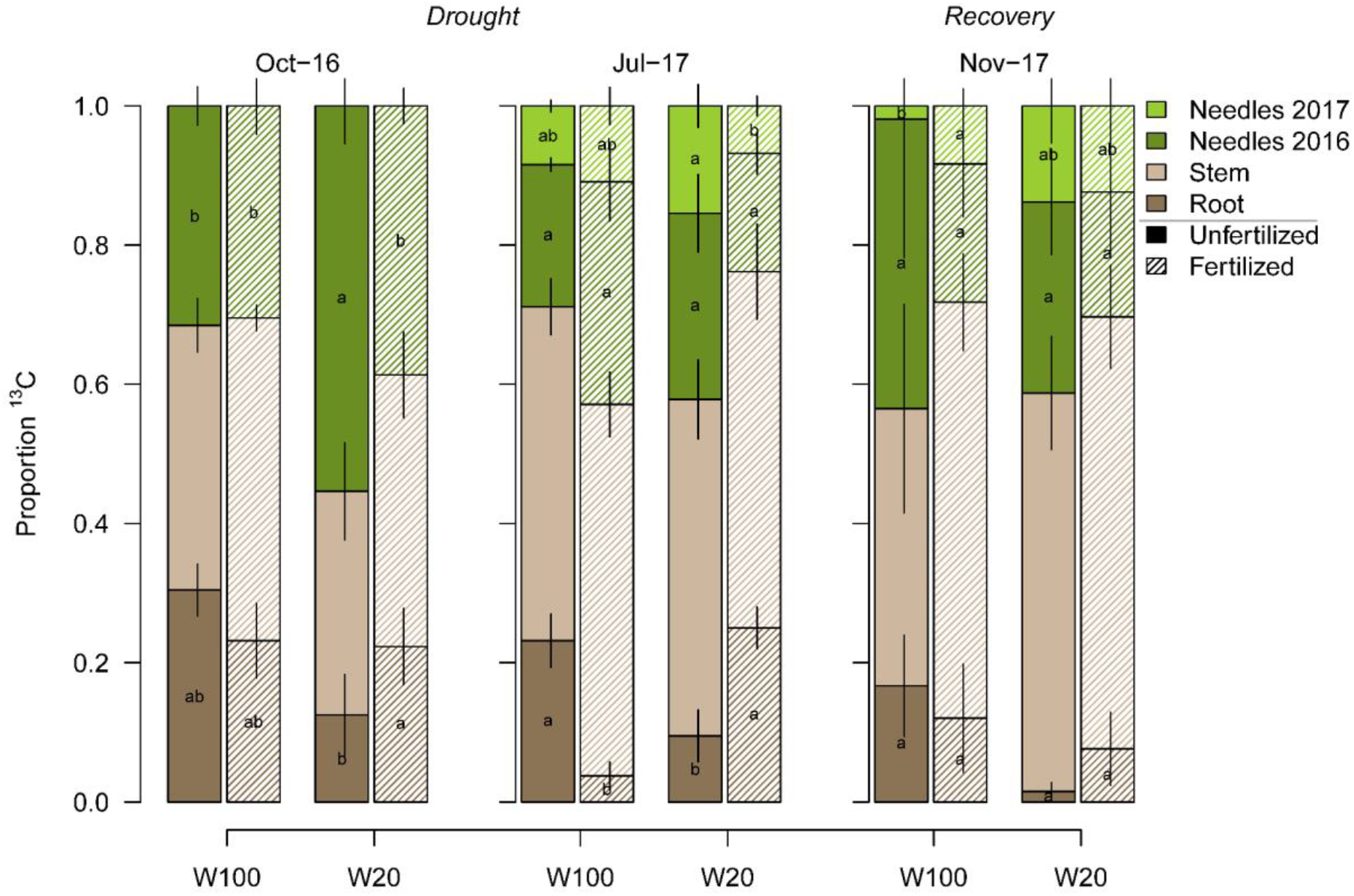
Proportion of total ^13^C found in the different tree compartments – needles produced in 2017 and 2016, stem and roots (indicated by colors). Solid bars indicate unfertilized, and bars with pattern indicate fertilized trees. Letters in the bars indicate significant differences between water and fertilization treatments within every tissue and harvest date. Error bars show the standard error of the mean (N=4).

### ^15^N incorporation and distribution

After the first year of extreme drought (W0) in October 2016, the ^15^N excess was significantly reduced in the stem and needles but only slightly (and not significant) in the roots, compared to well-watered W100 trees (Figure 3, Supplementary data Table S5), whilst W20 trees did not differ from W100. During the second drought year in July 2017, ^15^N excess in the needles and roots was much higher in W20 trees than in W100 trees. After rewetting, the ^15^N incorporation in previously W0 trees increased steeply in needles and stem, resulting in comparable amounts of ^15^N in all treatments, and decreased in the roots, resulting in lower amounts of ^15^N in the roots of W0 compared to W100 or W20 trees (Figure 3). The very high variance in the ^15^N excess in tissues of W0 trees were due to a lower number of replicates after high mortality events, and probably also due to high variation in recovery potential of previously drought stressed trees.

**Figure 3.**
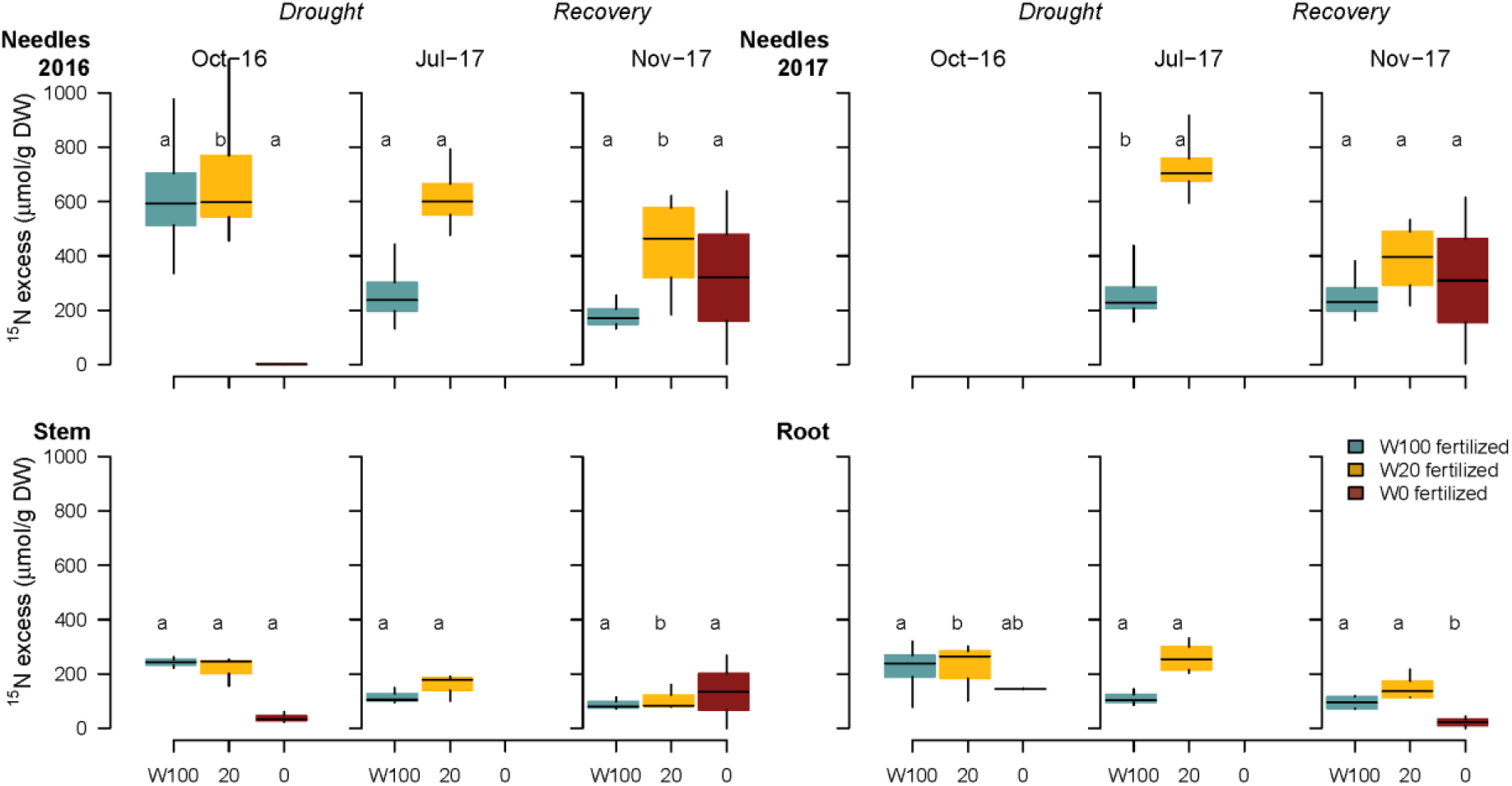
^15^N excess in needles produced in 2017 and 2016, stem and roots in October 2016, July 2017 and November 2017 (three months after rewetting). In July 2017, no samples were taken in the W0 treatment. Water regimes are indicated by colors, the shading indicates that only fertilized trees were tested. Letters indicate significant differences between water treatments within every tissue and harvest date. N=4 for every boxplot.

By October 2016, trees in W100 and W20 transported 57% (± 7%) and 70% (± 2%), respectively, of the total ^15^N taken up to their needles (ns between water treatments), whilst in W0 trees, the majority (72% ± 4%) of the ^15^N stayed in the roots (Figure 4). Only after rewetting, trees from the extreme drought treatment transported a significant amount of N towards needles. This caused similar distribution patterns in W0 trees compared to W100 and W20 trees (between 46% - 62% in needles and 9% - 14% in roots), with the exception of the newly grown needles, that received only a minor percentage of ^15^N. The proportion of ^15^N recovered in the stem was generally constant between harvest dates and water treatments (Figure 4).

**Figure 4.**
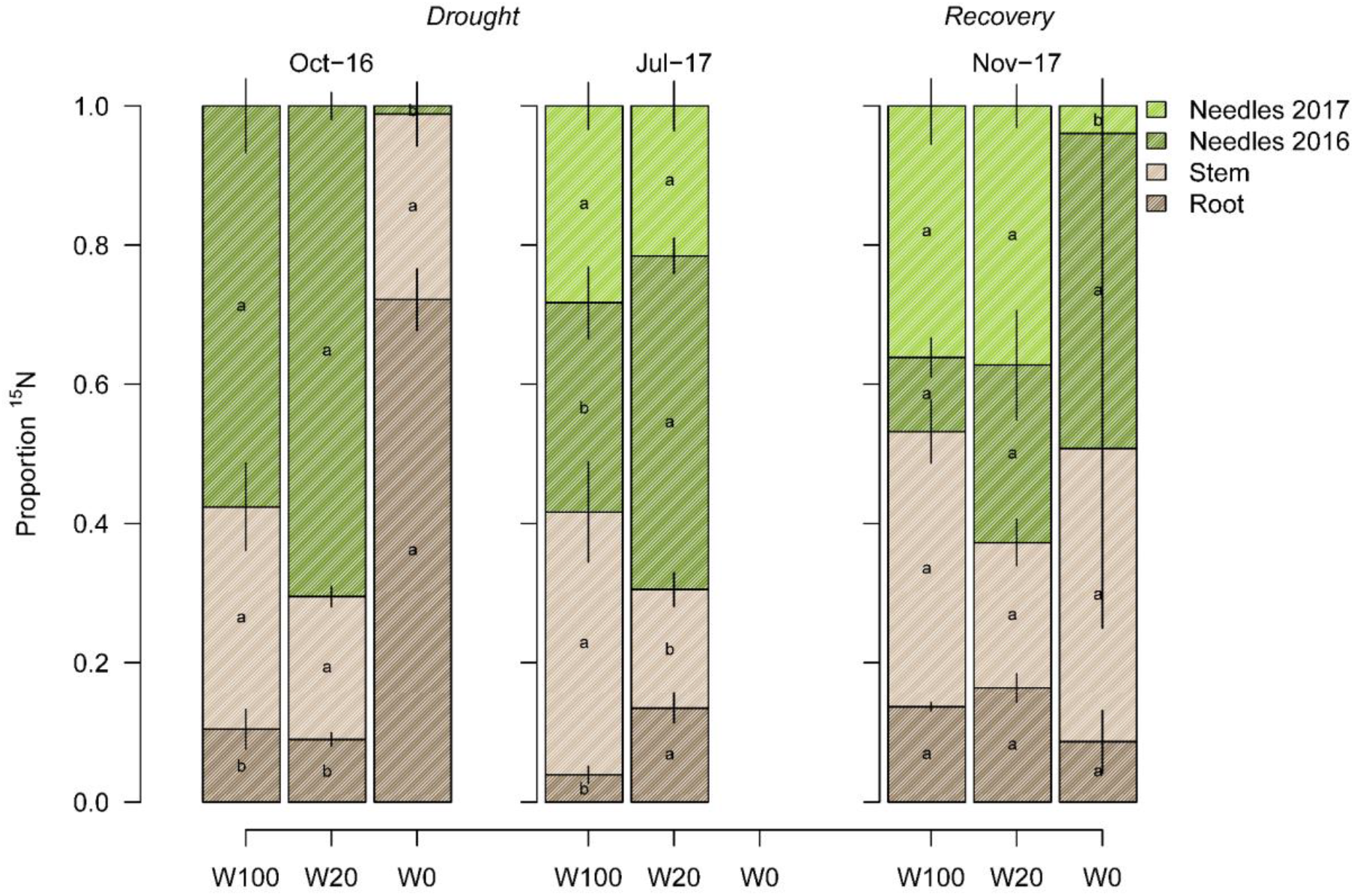
Proportion of total ^15^N found in the different tree compartments – needles produced in 2017 and 2016, stem and roots (indicated by colors). Pattern shows that only fertilized trees were labelled and measured. Letters indicate significant differences between water treatments within every tissue and harvest date. Error bars show the standard error of the mean (N=4).

## Discussion

We used a pulse-labelling ^13^C method and a ^15^N soil injection to assess the partitioning of C and N over 1.5 growing season. The experiment in the open-top chambers allowed us to only measure the background ^13^C and ^15^N once, before fumigation, and thus natural abundance of ^13^C is not known for later timepoints. However, we can assume that variation in natural abundance between the time of labelling and harvests is negligible. First of all, the variation in natural abundance was lower than the range of the measured label signal. Moreover, the trees were already drought stressed at time of labelling and the natural abundance of ^13^C in W0 (not fumigated with ^13^C) trees did not change during the two growing seasons, providing good reason to assume that this was also the case for W20 trees. In this study, we aimed for assessing the long-term distribution of label in structural and storage pools of the plants, while not capturing any respiration fluxes or leaf shedding. This prevents us from being able to integrate these measurements into a more closed mass balance (for example in Litton et al. 2007). Having said this, our approach allows for a long-term monitoring of single-pulse labelled C in trees, especially with regard to investment in newly grown tissues (next generation needles). Lastly, the injection of ^15^N was done with the most precise method available, but we acknowledge that the distribution and uptake of ^15^N in drought-prone plots (W20, W0) may still be limited due to limited mobility in the soil. We consider it important to focus mainly on the relative distribution of the labelled compound within the tree rather than looking at absolute values, to eliminate the limitations of this method.

### Interaction between water and nutrition drives changes in belowground C allocation

We hypothesized an increase of C allocation to the roots relative to other tissues under the W20 drought, to improve the water uptake potential (Kozlowski and Pallardy 2002, Freschet et al. 2018). However, under unfertilized conditions, the ^13^C allocation to roots was much lower in the drought treatment compared to well-watered trees, both in terms of ^13^C excess in roots and relative distribution within the plant (Figure 1, Figure 2). Hence, we had to reject our first hypothesis. Our previous assumption was that the W20 drought regime could be considered as a mild drought, because many other physiological parameters such as predawn leaf water potential, gas exchange, and biomass were only slightly but not significantly affected and no mortality occurred in the W20 drought regime (Supplementary data Table S3, S4) (Schönbeck et al. 2020). In contrast, the reduction in ^13^C allocation to roots suggests that soil water restriction might already have been severe enough to disable transport of new assimilates to the roots. One possibility would be that root biomass still increased with the use of older C reserves instead of newly assimilated C. Indeed, in our previous study, we found that C reserves such as starch and mobile sugars decreased with drought in the roots in October 2016. However, they were restored in July 2017 and total root biomass rather decreased in the W20 unfertilized treatment compared to W100 (Supplementary data Table S4) (Schönbeck et al. 2020). Alternatively, the metabolic activity of the roots might have been impaired by drought and thus C demand was restricted (Hagedorn et al. 2016). Considering that root embolisms are probably the first to occur during severe drought stress (Rodríguez-Calcerrada et al. 2017) and root NSCs are the most sensitive and variable compared to NSC in all other tissues (Hartmann et al. 2013, Choat et al. 2018), we can speculate that dysfunction and tree mortality is initiated in the root system during extreme drought stress.

We hypothesized that fertilization reduces assimilate allocation to roots compared to aboveground biomass in well-watered trees, but that fertilization in combination with drought increases C allocation belowground, due to the maintenance of root metabolism by improved nutrient uptake. Assimilate allocation to roots was slightly but not significantly reduced due to fertilization under well-watered conditions, which was also described in earlier studies (Kozlowski and Pallardy 2002, Gessler et al. 2017). Under non-limiting water conditions and increased nutrient availability, trees do invest more in aboveground biomass, causing lower root:shoot ratios. Under limiting water conditions, in accordance with our hypothesis, fertilization seemed to increase allocation of new assimilates to the roots. Nitrogen uptake of plants depends on the N availability to the roots, which is partially determined by the water mass flow and the nitrogen transported with it. Thus drought can, under constant soil nutritional conditions, cause nutrient limitation within plants (Figure 5). We speculate a positive feedback loop between drought and nutrient availability in the soil: increased nutrient availability in the soil improved the root nutrient uptake and released the nutrient limitation that was induced by drought (Figure 5). The higher nutrient uptake could then trigger plant responses to drought by stimulating e.g. the synthesis of drought-responsive amino acids and proteins (Alam et al. 2010). These compounds play a central role in osmoprotection (Nguyen and Lamant 1988, Rathinasabapathi 2000, Ashraf and Foolad 2007, Galiano Pérez et al. 2017) and might strengthen the C sink function of the roots. We thus expect that as a consequence of improved root activity and cell integrity, sink activity was increased as indicated by increased C allocation belowground (Figure 5).

**Figure 5.**
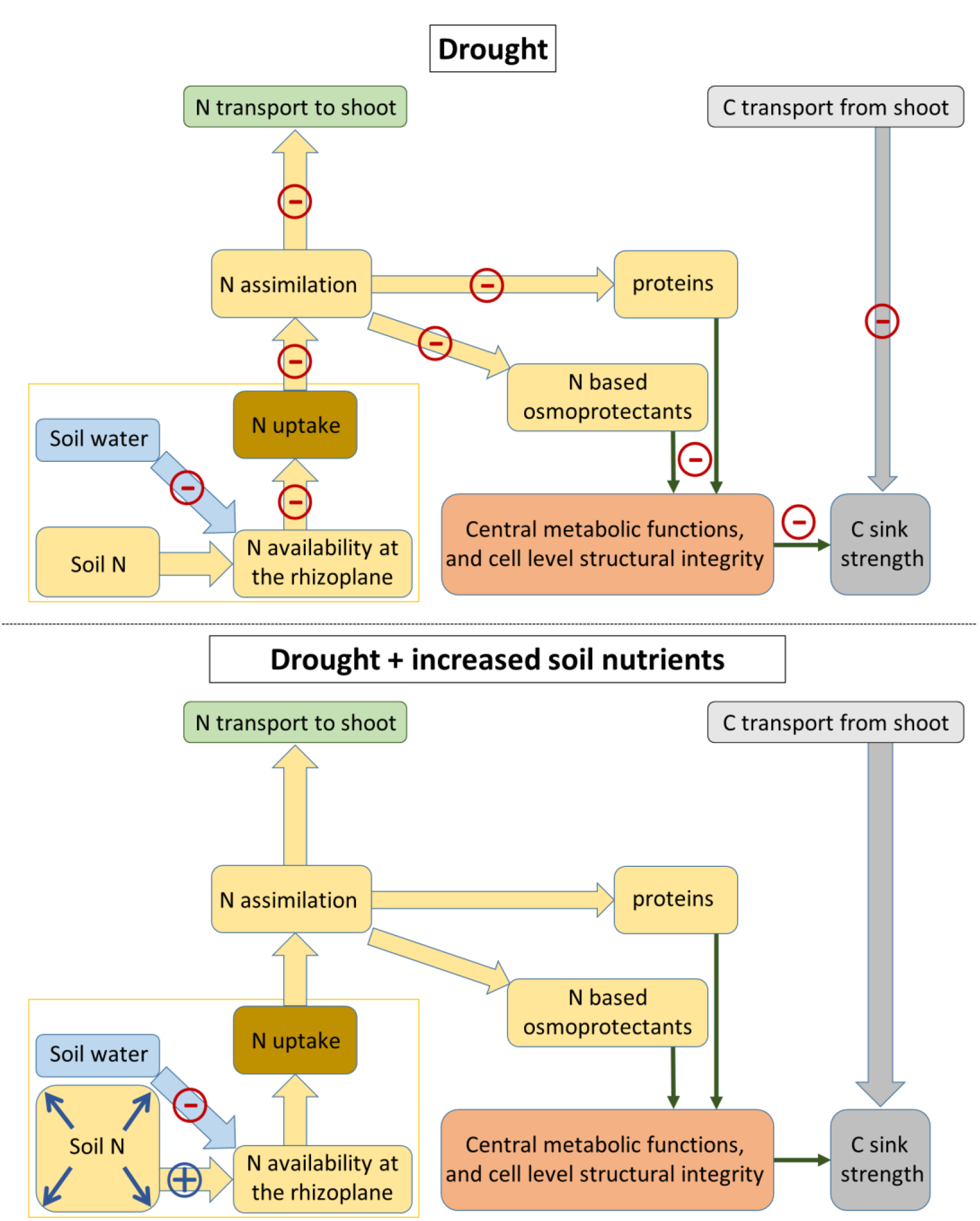
Conceptual framework on the role of soil N and drought in the allocation of assimilates. As both soil water availability (via water mass flow and thus transport of nitrogen) and soil nitrogen concentration influence the N availability at the rhizoplane, both can induce nutrient limitation to the plant. During drought, N based osmoprotectants might play an important role in maintaining central metabolic functions, sustaining or increasing the C sink strength and the C transport from the shoots to the roots. Drought might induce a nitrogen limitation because of decreasing transport of nutrients to the root surface (rhizoplane). An increase in soil N concentration could mitigate such N limitation due to reduced water mass flow and induce a positive feedback.

### Interaction effects between water and nutrition alters the C and N source of new needles

We hypothesized a strongly coordinated supply – demand regulation of C and N and thus expected that drought stressed trees do not use more stored C (i.e. ^13^C) and N (i.e. ^15^N) for growth of new needles than well-watered trees. The incorporation of ^13^C in new needles per dry weight was, however, higher in W20 than in W100 trees in July 2017 (Figure 1). The difference in turnover rate of ^13^C as well as a dilution of ^13^C due to higher needle biomass in W100 trees could have led to the differences found in the absolute values of incorporation. But when looking at the proportional distribution within the tree, it became clear that indeed W20 trees allocated relatively more ‘old’ C into new needles, at least when unfertilized. Fertilization cancelled out this allocation pattern and the opposite as in unfertilized trees was observed in reaction to drought. Although fertilization in well-watered conditions resulted in higher C allocation to the new needles, resulting in an increase of the aboveground biomass and photosynthetic active area as postulated by Gessler *et al*. (2017), fertilized trees under drought had the lowest relative amount of old C out of all treatments in the new needles. Considering nitrogen, we found that the ^15^N amount per g needle was much higher in W20 compared to W100, while the proportion of total ^15^N in the needles was similar between well-watered and mildly drought stressed (W20) trees, indicating that needle contribution to the total ^15^N pool in W20 trees was only small, likely caused by small needle biomass. These findings do not lead to an acceptance of our third hypothesis, especially regarding C in unfertilized conditions, probably due to transport failure. Moreover, ceasing of the root system in unfertilized drought-stressed trees might have increased the importance of needles (and stem) as a storage tissue. At the same time, spatial imbalances might have occurred (Klein et al. 2014), where root but not needle functionality was affected by drought, not inducing any stress related changes in prioritization of C into growth or storage. Furthermore, coniferous trees are thought to be less dependent on stored C for spring regrowth than deciduous trees – in our experiment, only 10% of ^13^C ended up in new grown needles, compared to levels up to 50% of stored C found in deciduous trees (Hansen and Beck 1990). Hence the risk and the consequences of maintaining or changing the relative amount of reallocated C to new-grown needles under drought are relatively low.

### Water availability after drought stress alters C and N allocation

For the recovery period, we expected that previously extreme drought stressed trees show an enhanced uptake and (re)allocation of N and a prioritization of C allocation belowground in response to rewetting, in order to restore the root system. Indeed, a shift was found in the allocation of ^15^N in previously extreme drought-stressed trees. Extreme drought (W0) initially inhibited ^15^N uptake by the roots and transport up to the needles (Figure 3) after the first year of drought and the little amount of N taken up was concentrated in the roots (Figure 4). On the one hand, ceasing of xylem and phloem transport probably influenced the N distribution between below- and aboveground tissues. Strongly reduced stomatal conductance (Schönbeck et al. 2020) indicates low xylem transport in the W0 treatment. On the other hand, N allocation to the roots during drought is important to support drought tolerance in the form of osmoprotective aminoacids, as was previously shown in beech (Fotelli et al. 2002). Rewetting recovered N transport to the needles, and the distribution of ^15^N was comparable between all drought regimes in November 2017. N transport from the roots to the shoot is important to restore the photosynthetic system and support aboveground metabolism and / or growth (Palacio et al. 2018). Moreover, rewetting caused extremely low ^13^C-label allocation to roots of previously drought stressed trees (Figure 1), and thus does not directly point to a prioritized C allocation belowground to restore the root system. We can thus not accept our last hypothesis regarding C allocation. However, as gas exchange in previously drought-stressed trees recovered (Supplementary data Table S3), the isotopic signal has likely been diluted by (non-labelled) new assimilates that have been allocated to regeneration of the root system. This is in agreement with findings of Hagedorn et al. (2016) of a strong prioritized transport of new rather than stored assimilates to the root system after drought release in beech.

### Conclusion

We could show that mainly the root system was affected by an interaction of drought and fertilization, while the expected alterations in C allocation to aboveground tissues such as newly formed needles could not be proven. We speculate that the root system might have already been impaired by the 80% reduction of water availability when no fertilization was applied, indicated by reduced C allocation to the root system during drought. We also speculate that a positive feedback loop might exist where fertilization improves the metabolism and functioning of the roots and might restore drought-induced alterations in C and N allocation, by contributing to the maintenance of cellular functions (e.g., via osmotic adjustment), consequently strengthening C sinks. Thus, an increased nutrient supply under drought does not only improve leaf metabolic functioning and cell structural integrity as suggested by Gessler et al. (2017) but might also be compensating for drought-induced loss of root functioning, thereby mitigating drought stress of trees. Our findings demonstrate both the importance of a strong functioning root system, and the difficulty in detecting early-warning signals for tree mortality if the mortality process starts belowground. As soil nutrients might play an important role in mitigating drought stress of trees, their potential should be more deeply investigated for different tree species. With this information available, tree species selection for climate smart forestry could be better adjusted to prevailing soil conditions.

## Data and Materials availability

### Supplementary data

Figure S1. Schematic overview and timeline of the experiment

Figure S2. Microclimate in the 16 Open top chambers

Figure S3. N content, NH_4_, NO_3_ and P concentrations in the soil.

Figure S4. C:N ratios in root, stem and needles of harvested trees.

Figure S5. Photo of trees in the open-top chambers

Table S1. Soil characteristics before start of the experiment.

Table S2. Natural abundance of C, N, ^15^N and ^13^C at the time of labelling

Table S3. Photosynthesis, stomatal conductance and predawn water potential at the time of labelling

Table S4. Biomass of roots, stem and needles at the three harvests

Table S5. ANOVA table of the linear mixed effect model testing ^13^C and ^15^N excess in the tissues against water and fertilization treatments.

### Funding

LS was supported by Swiss National Fund grant (31003A_157126/1) and AG acknowledges support by the Swiss National Fund (31003A_159866). MML was supported by the SNF Ambizione grant “TreeCarbo” (PZ00P2_179978).

## Acknowledgements

The authors thank Peter Bleuler for the technical organization of the open-top chamber facility, Matthias Haeni for the continuous data capturing, Kruno Sever and Rahel Mösch for their help with measurements and sampling, Jobin Joseph and Raphael Weber for their valuable assistance during labelling, Manuela Öttli and Matthias Saurer for conducting stable isotope measurements, Pierre Vollenweider, Jörg Luster and Frank Hagedorn for useful discussions on the methodology and treatments, and four anonymous reviewers for greatly improving the manuscript.

## Authors’ contributions

MHL, AG and AR conceived the study. All authors contributed equally to the planning and design of the experiment. LS, MHL, MML, MS and AG carried out the experiment. LS and MML collected the data. LS analyzed the data and all authors contributed to the interpretation of the results. LS constructed the first draft of the manuscript, all authors contributed significantly to the revisions and improvements of the manuscript.

