## Supplementary material for "Soil nutrient availability alters tree carbon allocation dynamics during drought"

*Tree Physiology*

**Figure S1:** Schematic overview of the 16 open-top chambers in a Latin square design. Blue indicates W100 treatment, Orange W20 and red W0. Solid colors are unfertilized lysimeters, and dashed areas are fertilized. Below a timeline of the years 2016 and 2017, where $^{15}N$, $^{13}C$ label and harvests are indicated. One water treatment (W50) was left out of this study, they are covered with grey hexagons.

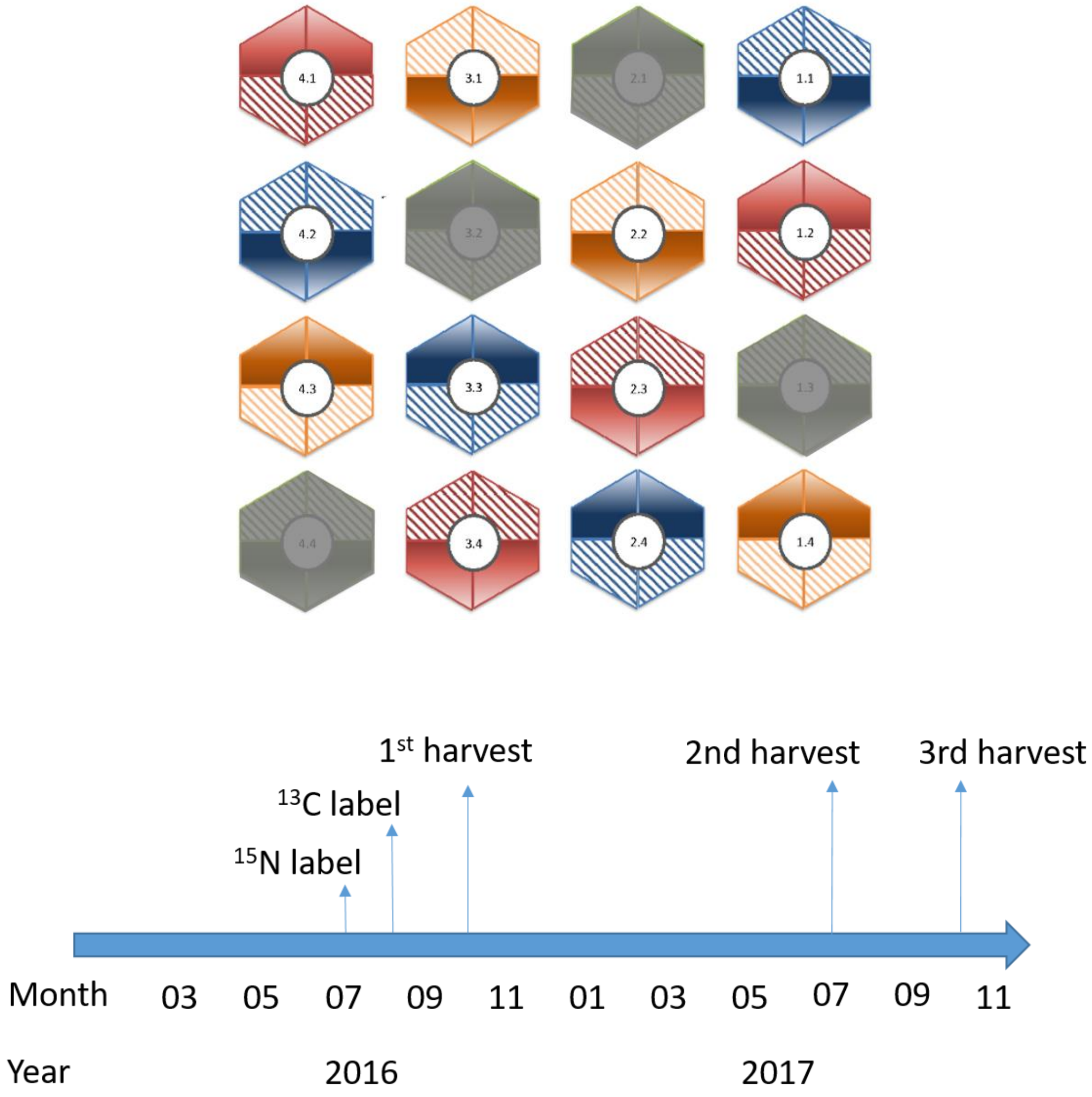

**Figure S2**. Air temperature, relative humidity and volumetric water content (VWC) in 20 cm soil depth in the four water treatments. VWC is separately shown for the unfertilized (solid lines) and fertilized (dashed lines) lysimeters. Vertical dashed lines in the VWC plot indicate the three harvests.

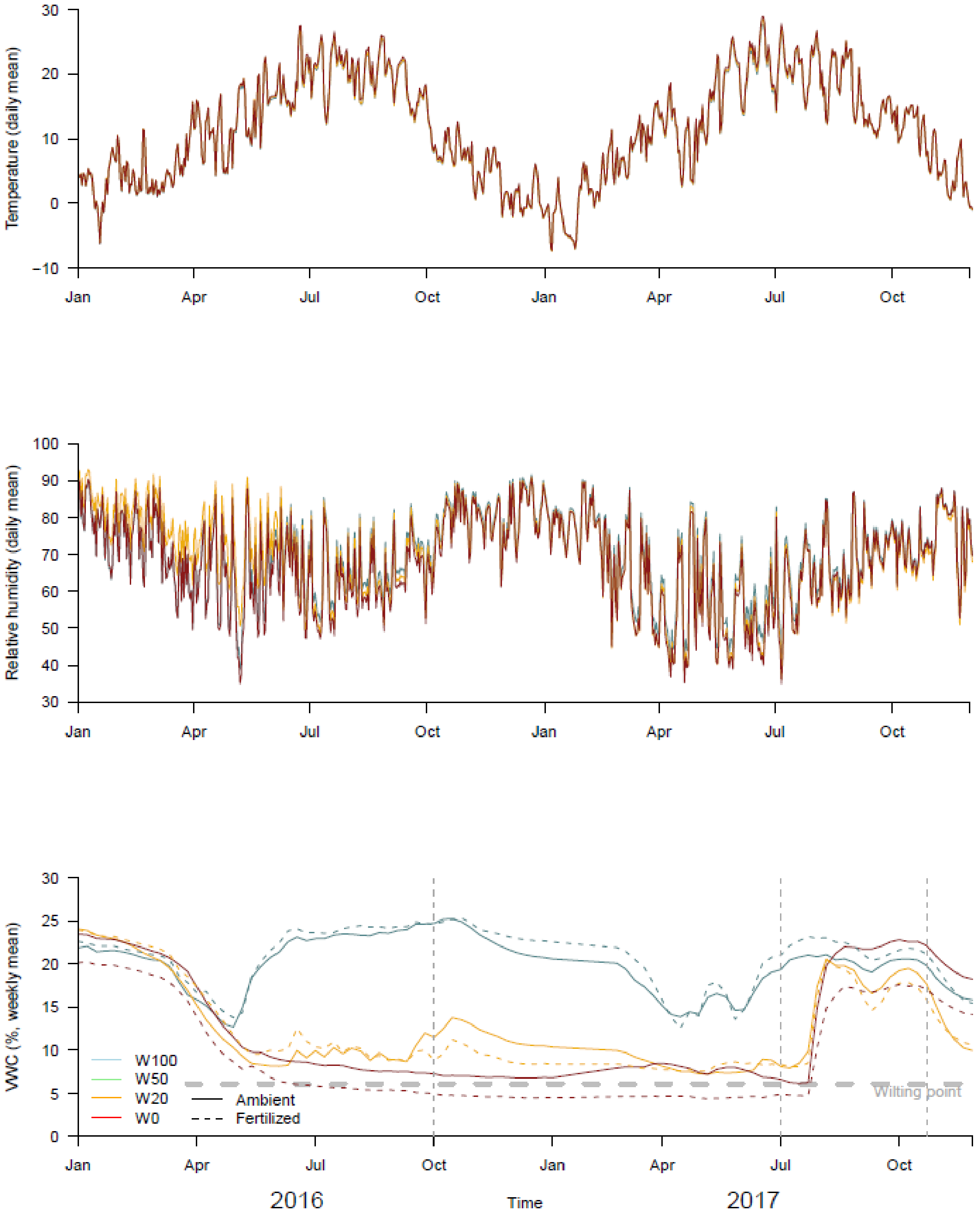

**Figure S3**. Total Nitrogen in grams in the ecosystem soil (colors) and plants (white bars) in October 2016, ammonium (NH4) concentration (mg/L), nitrate (NO3) concentration (µmol/g) and phosphorus (P) concentration (mg/g) in the soil. Colors represent the four drought treatments, W100 (blue), W20 (orange) and W0 (red). Patterns indicate fertilization treatments. Asterisks indicate significant differences between unfertilized and fertilized treatments within a water treatment. Error bars indicate the SE of the mean.

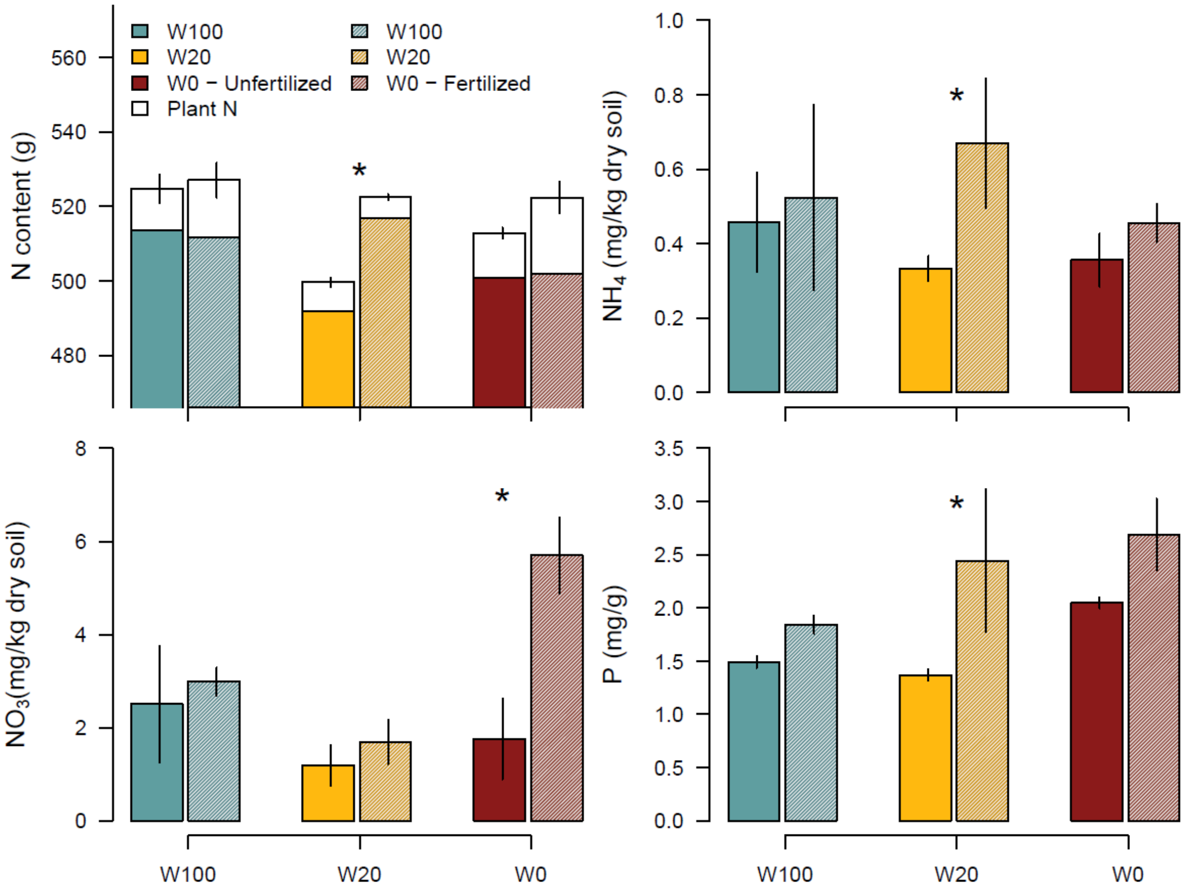

**Figure S4.** C:N ratios in needles produced in 2017 and 2016, stem and roots in October 2016, July 2017 and November 2017 (three months after rewetting). Water regimes are indicated by colors, fertilization is indicated by shading (solid = unfertilized, pattern = fertilized). Letters indicate significant differences between water and fertilization treatment within every tissue and harvest date, as tested by a mixed effect linear model. Error bars indicate standard error of the mean (SE).

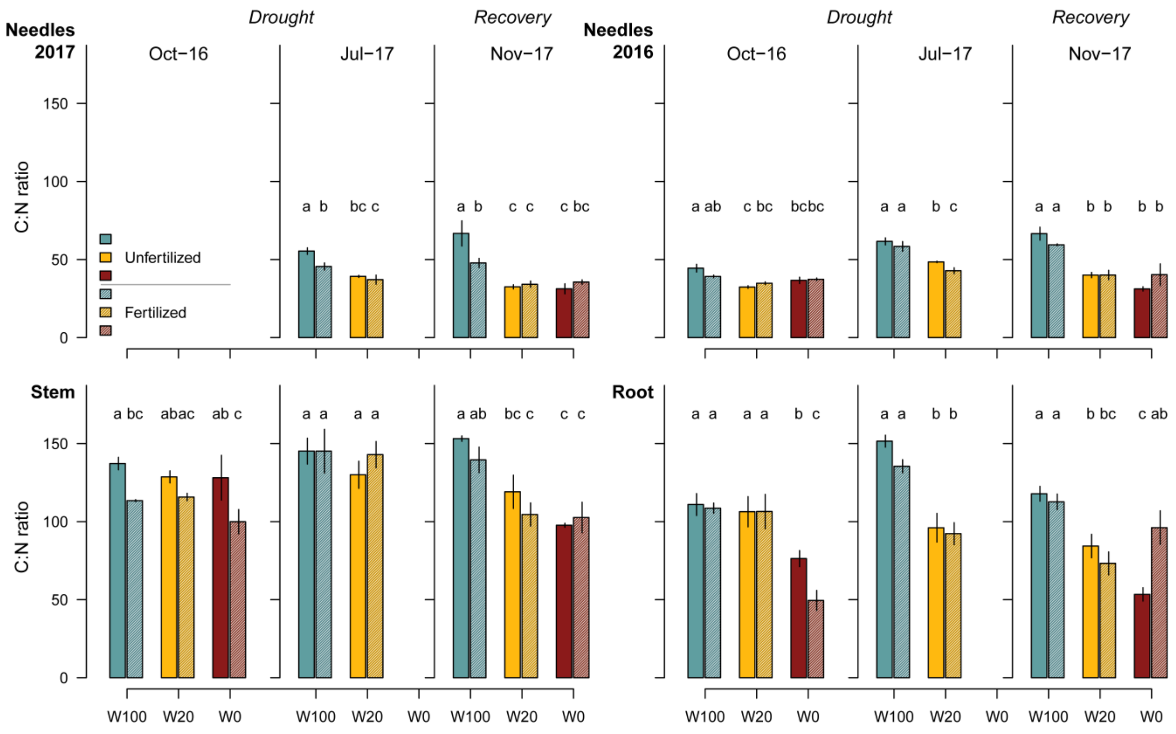

**Figure S5: P**hotograph of all chambers taken in May 2017 from the birds-eye perspective.

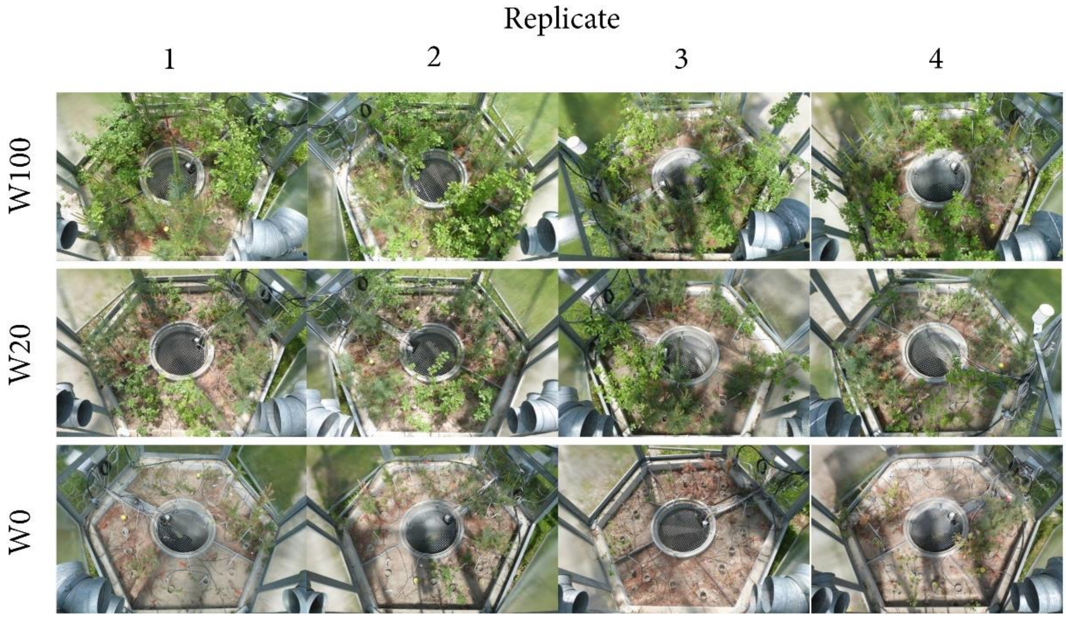

**Table S1:** Original soil characteristics before start of treatment in the open top chambers (OTCs)

| Characteristics | Calcareous sandy-loam |
|---|---|
| Origin | Brugg (Fluvisol) |
| Texture (% sand, silt, clay) | 71, 18, 12 |
| pH (0.01 $_{M}$ $CaCl_2$) | 6.88 |
| $C_{tot}$ (%) | 1.97 |
| $N_{tot}$ (%) | 0.05 |
| $P_{tot}$ ($mg \cdot kg^{-1}$) | 357.96 |
| $Ca_{exch.}$ ($mg \cdot kg^{-1}$) | 1629.46 |
| $Mg_{exch.}$ ($mg \cdot kg^{-1}$) | 21.87 |
| $K_{exch.}$ ($mg \cdot kg^{-1}$) | 25.75 |
| $Mn_{exch.}$ ($mg \cdot kg^{-1}$) | 1.44 |
| CEC ($mmol_c \cdot kg^{-1}$) | 84.69 |
| Base saturation (%) | 99.45 |

**Table S2:** Natural abundances of C, N, $\delta^{13}C$ and $\delta^{15}N$ at the time of labelling (July 2016). Letters indicate significant differences between treatments as tested by mixed effect linear models.

***Needles***

| **Treatment** | **C** | **N** | **$d^{13}C$** | **$d^{15}N$** |
|---|---|---|---|---|
| ***W100 – A*** | $46.58 \pm 0.13^{a}$ | $0.88 \pm 0.12^{a}$ | $-30.26 \pm 0.09^{a}$ | $4.34 \pm 0.53^{a}$ |
| ***W100 – N*** | $46.4 \pm 0.16^{a}$ | $0.98 \pm 0.11^{a}$ | $-30.37 \pm 0.29^{a}$ | |
| ***W20 – A*** | $47.69 \pm 0.7^{b}$ | $1.03 \pm 0.12^{a}$ | $-29.34 \pm 0.24^{b}$ | $4.89 \pm 0.52^{a}$ |
| ***W20 – N*** | $46.79 \pm 0.26^{ab}$ | $1.14 \pm 0.16^{a}$ | $-28.49 \pm 1.16^{b}$ | |

***Stem***

| **Treatment** | **C** | **N** | **$d^{13}C$** | **$d^{15}N$** |
|---|---|---|---|---|
| ***W100 – A*** | $46.35 \pm 0.73^{a}$ | $0.37 \pm 0.04^{a}$ | $-28.08 \pm 0.33^{a}$ | $33.22 \pm 9.78^{a}$ |
| ***W100 – N*** | $45.99 \pm 0.48^{b}$ | $0.43 \pm 0.06^{a}$ | $-28 \pm 0.71^{a}$ | |
| ***W20 – A*** | $46.34 \pm 0.81^{a}$ | $0.39 \pm 0.04^{a}$ | $-26.24 \pm 1.78^{b}$ | $32.89 \pm 3.52^{a}$ |
| ***W20 – N*** | $45.91 \pm 0.66^{b}$ | $0.44 \pm 0.03^{a}$ | $-25.69 \pm 2.27^{b}$ | |

***Roots***

| **Treatment** | **C** | **N** | **$d^{13}C$** | **$d^{15}N$** |
|---|---|---|---|---|
| ***W100 – A*** | $43.8 \pm 3.22^{a}$ | $0.41 \pm 0.05^{a}$ | $-30.76 \pm 0.09^{a}$ | $10.52 \pm 2.06^{a}$ |
| ***W100 – N*** | $46.92 \pm 6.4^{a}$ | $0.45 \pm 0.05^{a}$ | $-30.97 \pm 0.29^{a}$ | |
| ***W20 – A*** | $47.07 \pm 3.64^{a}$ | $0.47 \pm 0.06^{a}$ | $-29.34 \pm 0.24^{b}$ | $10.17 \pm 2.65^{a}$ |
| ***W20 – N*** | $45.93 \pm 4.15^{a}$ | $0.58 \pm 0.22^{a}$ | $-28.49 \pm 1.16^{b}$ | |

**Table S3:** Photosynthetic activity, stomatal conductance, and predawn needle water potential in the three water treatments and two fertilization treatments before the three harvests. Values indicate mean +/- SE. Letters indicate significant differences between water x fertilization treatments as tested by mixed effect linear models.

| Water | Fertilization | Photosynthesis (μmol/m²/s¹) | Stomatal conductance (mol/m²/s¹) | Predawn water potential (Ψ) |
|---|---|---|---|---|
| *October 2016* | | | | |
| **W100** | A | 9.07 ± 0.64[a] | 0.206 ± 0.017[a] | 0.24[a] |
| | N | 12.07 ± 1.07[a] | 0.249 ± 0.020[a] | 0.32[a] |
| **W20** | A | 6.36 ± 1.00[a] | 0.123 ± 0.031[b] | 0.63[a] |
| | N | 6.25 ± 0.79[a] | 0.063 ± 0.016[b] | 0.57[a] |
| **W0** | A | 0.99 ± 0.35[b] | 0.012 ± 0.006[b] | 0.38[b] |
| | N | 1.11 ± 0.38[b] | 0.012 ± 0.010[b] | 0.43[b] |
| *July 2017* | | | | |
| **W100** | A | 9.59 ± 0.62[a] | 0.374 ± 0.006[a] | 0.21[a] |
| | N | 7.44 ± 0.71[a] | 0.203 ± 0.051[a] | 0.16[a] |
| **W20** | A | 8.80 ± 1.21[a] | 0.165 ± 0.029[a] | 0.47[a] |
| | N | 5.6 ± 0.88[a] | 0.063 ± 0.018[a] | 0.37[a] |
| **W0** | A | 1.19 ± 0.51[b] | 0.004 ± 0.003[b] | 1.18[b] |
| | N | 0.00 ± 0.06[b] | 0.002 ± 0.000[b] | 1.10[b] |
| *October 2017* | | | | |
| **W100** | A | 9.10 ± 0.78[a] | 0.163 ± 0.009[a] | 0.35[a] |
| | N | 7.65 ± 0.51[a] | 0.157 ± 0.008[a] | 0.35[a] |
| **W20** | A | 7.51 ± 0.54[a] | 0.122 ± 0.012[a] | 0.30[a] |
| | N | 8.70 ± 0.46[a] | 0.117 ± 0.009[a] | 0.42[a] |
| **W0** | A | 10.48 ± 1.11[a] | 0.173 ± 0.025[a] | 0.50[a] |
| | N | 5.65 ± 0.85[a] | 0.069 ± 0.001[a] | 1.07[a] |

**Table S4:** Biomass of roots, stem and needles at the harvest dates (in grams). Values show mean +/- SE. Letters in superscript indicate significant differences between water treatments. A '+' in superscript means a significant fertilization effect. Tested with linear mixed effect model.

| | | Root | Stem | Needles |
|---|---|---|---|---|
| **Water** | **Fertilization** | | | |
| *October 2016* | | | | |
| W100 | A | 31.1 ± 4.9$^{a}$ | 63.0 ± 7.3$^{a}$ | 44.6 ± 5.8$^{a}$ |
| | N | 23.5 ± 3.1$^{a}$ | 52.8 ± 4.2$^{a}$ | 50.8 ± 8.8$^{a}$ |
| W20 | A | 15.8 ± 1.5$^{a}$ | 33.8 ± 3.2$^{a}$ | 26.8 ± 3.2$^{a}$ |
| | N | 28.5 ± 8.2$^{a}$ | 62.4 ± 8.2$^{a+}$ | 61.7 ± 7.8$^{a+}$ |
| W0 | A | 12.1 ± 1.3$^{b}$ | 32.3 ± 1.5$^{b}$ | 20.7 ± 2.2$^{b}$ |
| | N | 9.0 ± 1.4$^{b}$ | 21.7 ± 1.7$^{b}$ | 13.6 ± 1.1$^{b}$ |
| *July 2017* | | | | |
| W100 | A | 19.8 ± 2.4$^{a}$ | 70.4 ± 6.9$^{a}$ | 58.1 ± 8.1$^{a}$ |
| | N | 26.0 ± 6.1$^{a}$ | 115.1 ± 16.8$^{a}$ | 93.8 ± 9.6$^{a}$ |
| W20 | A | 14.9 ± 1.4$^{a}$ | 57.0 ± 5.3$^{a}$ | 53.0 ± 3.7$^{a}$ |
| | N | 32.9 ± 5.3$^{a}$ | 71.4 ± 7.4$^{a}$ | 72.9 ± 10.7$^{a+}$ |
| *November 2017* | | | | |
| W100 | A | 46.2 ± 4.2$^{a}$ | 200.9 ± 15.4$^{a}$ | 111.4 ± 12.0$^{a}$ |
| | N | 52.3 ± 5.3$^{a}$ | 127.5 ± 10.2$^{a}$ | 87.2 ± 3.9$^{a}$ |
| W20 | A | 50.6 ± 2.5$^{a}$ | 96.1 ± 5.6$^{b}$ | 87.3 ± 5.2$^{a}$ |
| | N | 51.8 ± 6.5$^{a}$ | 91.9 ± 8.8$^{b}$ | 80.6 ± 11.7$^{a}$ |
| W0 | A | 16.9 ± 2.5$^{b}$ | 23.3 ± 3.4$^{c}$ | 17.4 ± 1.4$^{b}$ |
| | N | 18.2 ± 1.5$^{b}$ | 44.6 ± 5.5$^{c}$ | 14.0 ± 2.8$^{b}$ |

**Table S5.** Anova table of the linear mixed effect model testing $^{13}C$ and $^{15}N$ excess in the different tissues against water and fertilization treatment and their interaction. Separate models were made per harvest date. Values written in bold indicate significant effects. Pairwise comparisons can be found in figure 1 and 3 in the main text.

| | | **Root** | | **Stem** | | **N16** | | **N17** | |
|---|---|---|---|---|---|---|---|---|---|
| $^{13}$**C** | ***Df*** | ***F*** | ***p*** | ***F*** | ***p*** | ***F*** | ***p*** | ***F*** | ***p*** |
| ***October*** | | | | | | | | | |
| *Water* | 1 | 3.50 | 0.124 | 1.75 | 0.213 | 0.45 | 0.526 | | |
| *Fertilization* | 1 | 0.79 | 0.418 | 2.25 | 0.162 | 1.52 | 0.264 | | |
| *W:F* | 1 | 1.85 | 0.24 | 0.01 | 0.914 | 1.52 | 0.264 | | |
| ***July*** | | | | | | | | | |
| *Water* | 1 | 13.35 | **0.004** | 19.12 | **0.008** | 0.12 | 0.741 | 32.02 | **0.001** |
| *Fertilization* | 1 | 4.46 | 0.058 | 11.33 | **0.022** | 2.53 | 0.163 | 0.02 | 0.901 |
| *W:F* | 1 | 30.12 | <**0.001** | 8.79 | **0.034** | 1.97 | 0.210 | 0.93 | 0.373 |
| ***November*** | | | | | | | | | |
| *Water* | 1 | 4.36 | 0.082 | 1.99 | 0.186 | 0.00 | 0.988 | 1.35 | 0.29 |
| *Fertilization* | 1 | 0.19 | 0.679 | 0.56 | 0.470 | 3.05 | 0.131 | 0.54 | 0.488 |
| *W:F* | 1 | 1.37 | 0.286 | 0.00 | 0.966 | 0.57 | 0.480 | 2.42 | 0.171 |
| $^{15}$**N** | | | | | | | | | |
| ***October*** | 2 | 0.24 | 0.795 | 24.68 | **0.003** | 207.04 | <**0.001** | | |
| ***July*** | 1 | 21.20 | **0.006** | 1.29 | 0.320 | 11.94 | **0.014** | 21.84 | **0.003** |
| ***November*** | 2 | 6.01 | **0.030** | 0.73 | 0.529 | 1.46 | 0.296 | 1.39 | 0.309 |
